# Functional Interrogation of Neuronal Subtypes via Intersectional Expression of Optogenetic Actuator Reveals Non-linear Components in a Linear Circuit

**DOI:** 10.1101/2025.02.03.636179

**Authors:** Tiannuo Li, Sandeep Kumar, Hoikiu Poon, Andrew M Leifer, Chaogu Zheng

**Affiliations:** School of Biological Sciences, The University of Hong Kong, Hong Kong SAR, China; Princeton Neuroscience Institute, Princeton University, Princeton, New Jersey, United States of America; Department of Physics, Princeton University, Princeton, New Jersey, United States of America

**Author notes:** These authors contributed equally to the study (Tiannuo Li, Sandeep Kumar). Correspondence (C.Z.) and (A.M.L.).

## Abstract

Investigating signal integration in a neural circuit is oftentimes challenging when the circuit contains neuronal subtypes that are transcriptomically similar, due to the lack of tools to express optogenetic actuators with high cellular specificity and to deliver light with high spatiotemporal accuracy. Here, we demonstrate the use of a split GAL4-based genetic “AND” gate to express Chrimson in specific touch receptor neuron (TRN) subtypes in the *C. elegans* touch response circuit. Combining this intersectional strategy for transgene expression with high-throughput optical targeting and behavioral quantification, we optogenetically interrogated the role of each TRN subtype in mediating the mechanosensor-induced escape response and in integrating signals that trigger the opposite motor output. Surprisingly, we found that although the response of the overall circuit linearly combines the competing anterior and posterior stimuli, this linearity is comprised of antagonistic non-linear contributions from the anterior and posterior sensors, which conspire to generate a linear response.

## Introduction

Understanding how opposing signals are integrated in a neural circuit is a fundamental but challenging task in neuroscience. In theory, targeted activation through the expression of an optogenetic actuator (e.g., channelrhodopsin)^1,2^ in specific neurons of a circuit can enable the functional assessment of the neuron’s contribution to the circuit, and one could investigate signal integration by simultaneously activating neurons that trigger the opposite behavioral responses in the same circuit. This approach, however, is often technically difficult due to the lack of cell-specific promoters. One potential solution is to use an intersectional strategy to drive gene expression in cells where the expression patterns of two distinct promoters overlap through genetic “AND” gates,^3,4^ such as the split GAL4,^5,6^ split Q,^7^ and split Cre systems.^8^ The common principle of these approaches is to split a gene expression driver into two non-functional modular halves, which can then reconstitute a functional protein (when expressed in the same cell) to activate effector expression. Although these methods were proven in principle, examples of using the intersectional strategy to discern the function of closely related neuronal subtypes are still rare.

In this study, we applied a previously established split GAL4 system^6^ to label and activate specific subtypes of the mechanosensory touch receptor neurons (TRNs) and assessed their roles in integrating antagonizing signals in the touch response circuit in *Caenorhabditis elegans*. The six TRNs (ALML/R, AVM, PLML/R, and PVM) detect gentle body touch and transduce mechanical stimuli into neuronal signals,^9^ which are relayed to downstream interneurons to trigger either forward or backward movement in avoidance response (Figure 1A-B). Among the four subtypes, ALM and PLM have two bilaterally symmetric left and right neurons, while AVM and PVM have only one neuron. All six neurons express largely the same genetic program associated with the TRN fate,^10,11^ although subtle differences in their transcriptomes were found.^12^ Previous cell ablation experiments found that ablating ALML/R or AVM neurons caused adult animals to be partially touch-insensitive, while ablating all three neurons led to a complete loss of anterior sensitivity.^9^ Ablation of PLML/R led to the loss of posterior sensitivity, whereas elimination of PVM had no effects on touch response.^9^ These studies revealed the relative importance of each TRN subtype in the touch response circuit, but it remains unclear how the activation of a single TRN subtype affects behavioral output. Optogenetic activation of individual TRN subtype is challenging due to the lack of promoters that are solely expressed in one subtype.

**Figure 1.**
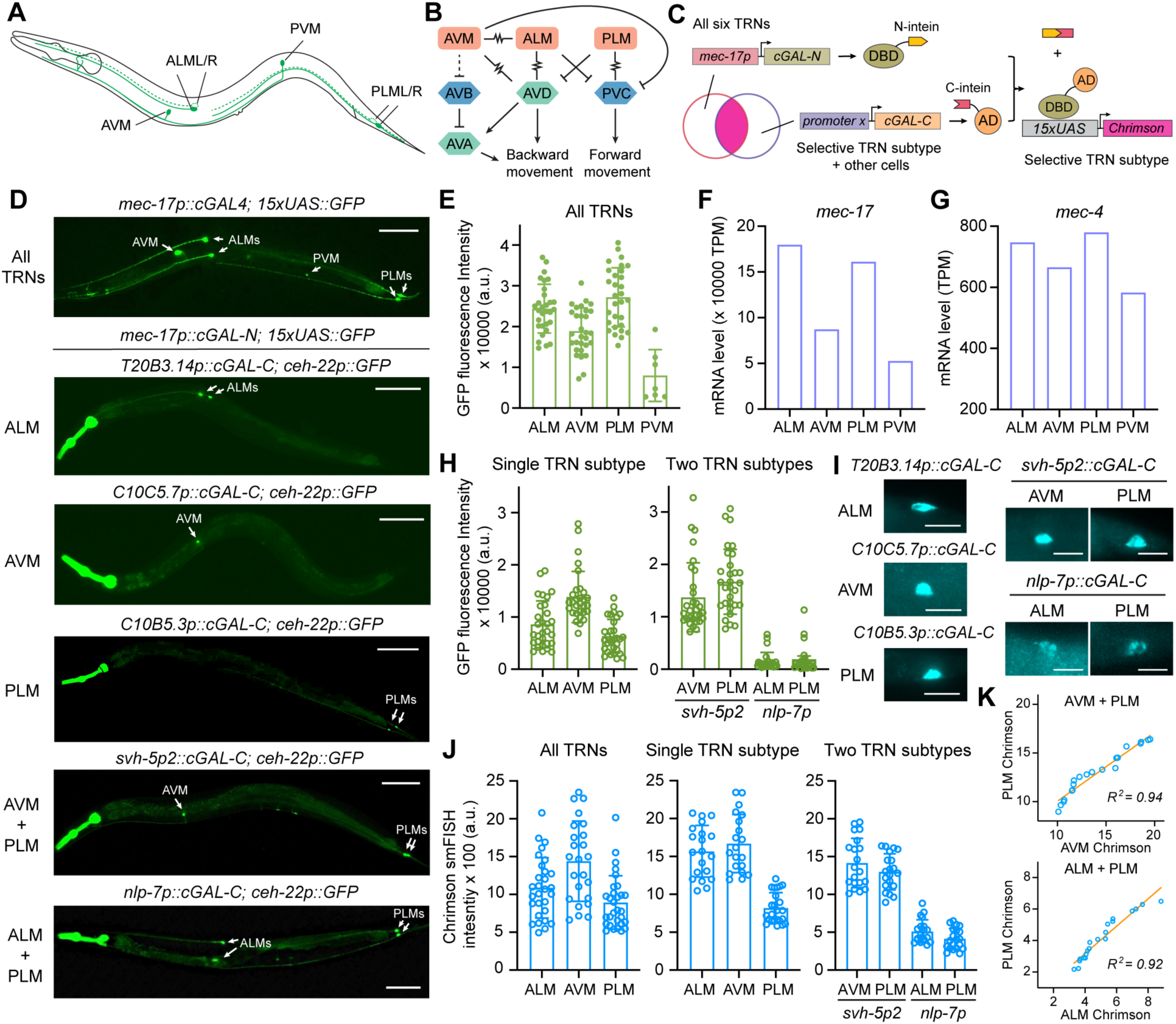
Expression of Chrimson in selective TRN subtypes. (A) Location and morphology of the six TRN subtypes. (B) Neuronal connection in touch circuit. Lines with zig zag indicate gap junction connection. Other lines indicate either excitatory or inhibitory chemical synapses. (C) Strategies for intersectional expression using the *mec-17* promoter to drive the expression of cGAL-N in the six TRNs and other candidate promoters (promoter x) to drive cGAL-C. The reconstituted cGAL binds to UAS and activates the expression of Chrimson. (D) Expression of GFP in animals carrying *syIs300[15xUAS::GFP]* and *unkIs17[mec-17p::cGAL-N]* and other transgenes that expressed cGAL-C from various promoters; *ceh-22p::GFP* was used as a co-injection marker and was expressed in the pharynx. Scale bars = 100 μm. (E) GFP intensity in the TRN subtypes of CGZ2180 *syIs595[mec-17p::cGAL4] V; syIs390[15xUAS::GFP] III* animals that express GFP in all TRNs. (F-G) The abundance of *mec-17* and *mec-4* transcripts (in Transcripts Per Million; TPM) in TRN subtypes according to previous scRNA-seq data.^17^ (H) GFP fluorescence intensity in the TRNs of animals that specifically label one or two TRN subtypes. (I) Signal of smFISH against Chrimson in the TRNs of animals that specifically express Chrimson in one or two TRN subtypes. Scale bars = 10 μm. (J) Quantification of the smFISH signal for Chrimson. (K) Correlation of the Chrimson smFISH signal in the two TRN subtypes in the same animal of the AVM-PLM and the ALM-PLM strains. Error bars in this Figure indicate standard deviation.

In certain experimental setup, precise optical targeting could achieve optogenetic activation of a single TRN subtype,^13^ but the approach is laborious and has low throughput. Targeted illumination allows the activation of anterior and posterior TRNs in free moving animals with higher throughput^14^ but could not achieve subtype specificity, since head illumination activate both ALM and AVM neurons and posterior illumination activates both PLM and PVM neurons. The lack of subtype-level specificity makes it difficult to understand the integration of opposing signals from two TRN subtypes (e.g., ALM and PLM) in the circuit. Thus, to stimulate only the desired TRN subtypes with throughput high enough for behavioral analysis, cell-specific genetic drivers are needed.

We developed such drivers by expressing two halves of the split cGAL (*C. elegans*-optimized GAL-4^6^) from two distinct promoters that only overlap in one TRN subtype (Figure 1C). Crossing these drivers with effectors that express Chrimson from a UAS promoter allowed us to activate individual TRN subtype and compare their circuit output in high-throughput behavioral analyses. Furthermore, by applying the targeted illumination on strains that express Chrimson in specific pairs of TRNs in the anterior and posterior halves of the animal, we investigated how mechanosensory signals from the subtypes that drive opposite response are integrated in the circuit and identified an unexpected non-linear component of the touch circuit. Overall, our study serves as an example of combining advanced genetic tools with high-throughput population-scale optogenetic manipulation and behavioral quantification to understand how neural circuits drive behavior.

## Results

### Screening promoters for intersectional expression in TRN subtypes

Our strategy was to use a TRN-specific promoter to express one half of the split cGAL in all six TRNs and then use another promoter that is differentially expressed among the TRN subtypes to drive the second half of the cGAL. This approach allows the reconstitution of cGAL only in some TRN subtypes but not the others. We first constructed a strain expressing the N-terminal DNA-binding domain of cGAL under the *mec-17* promoter, which is known to drive strong and exclusive expression only in the six TRNs.^11,15^ We then crossed the integrated *mec-17p::cGAL-N* transgene (*unkIs17*) with the effector *15xUAS::gfp* (*syIs300*).^16^ As expected, the cGAL-N half alone was not able to drive GFP expression in the TRNs. Next, we identified 18 genes that were expressed in only one or two TRN subtypes among the four subtypes based on single-cell transcriptomic data^17^ or previous fluorescent reporter studies.^12,18–20^ We used their promoters to express the C-terminal transcriptional activation domain of cGAL in animals that carried the *mec-17p::cGAL-N* and *15xUAS::gfp* transgenes. Among the 19 promoters we tested (two for *svh-5*), four promoters drove cGAL-C expression that led to GFP signals in only one TRN subtype, and three gave rise to GFP expression in two TRN subtypes (Table S1). Based on the intensity and stability of the GFP signals, we chose to use *T20B3.14*, *C10C5.7*, and *C10B5.3* promoters to drive cGAL-C expression in ALM, AVM, and PLM neurons, respectively, *nlp-7* promoter for expression in ALM and PLM neurons, and *svh-5a* promoter (named *svh-5p2*) for AVM and PLM expression (Figure 1D). Since PVM does not make synapses with the interneurons that control locomotion and ablating PVM did not have any detectable effects on touch sensitivity^9,21^, we did not focus on labeling PVM neurons specifically in this study.

We then integrated the cGAL-C-expressing transgenes into the genome of animals carrying both *unkIs7[mec-17p::cGAL-N]* and *syIs503[15xUAS::chrimson]* for TRN subtype-specific expression of Chrimson. To assess the expression level of Chrimson, we measured the activation of the 15xUAS promoter in each strain by crossing the integrated cGAL-C transgenes back into the *unkIs17; syIs300* animals to quantify GFP intensity. As controls, we first measured GFP expression in each TRN subtype in the *syIs595[mec-17p::cGAL4]; syIs300[15xUAS::GFP]* animals and found that GFP was expressed most strongly in ALM and PLM neurons; the expression was weaker in AVM and the weakest in PVM neurons (Figure 1E). This expression pattern is consistent with the *mec-17* mRNA levels among the four TRN subtypes, thus reflecting the varying activities of *mec-17* promoter in these subtypes (Figure 1F). The general trend also matched the abundance of *mec-4* mRNA, which codes for the mechanotransducer channel in the TRNs (Figure 1G), suggesting that ALM and PLM may be more mechanosensitive than AVM and PVM. This feature is recapitulated in the optogenetic strain, which has stronger UAS activation in ALM and PLM compared to the other two subtypes.

For the three strains with reconstituted cGAL in only one TRN subtype, GFP expression level is similar across ALM, AVM, and PLM neurons (Figure 1H). For the two strains expressing reconstituted cGAL in two TRN subtypes, GFP expression was found similar between the subtypes in the same strain, although the strain expressing cGAL in ALM and PLM neurons had weaker GFP expression than the strain expressing cGAL in AVM and PLM neurons (Figure 1H). Nonetheless, the transgene *unkIs54[nlp-7p::cGAL-C]* for ALM and PLM was on the same chromosome as *syIs300[15xUAS::GFP]*, which prevented us from replacing the Chrimson effector with the GFP effector. So, we used the extrachromosomal array to perform the GFP quantifications. We also tracked GFP expression across developmental stages in the above strains and found that expression in ALM and PLM could be observed from the first larval stage, while stable expression in AVM was only observed from the third larval stage (Figure S1), which matches the embryonic differentiation of ALM and PLM and the postembryonic development of AVM neuron.

To circumvent the problem of transgene linkage, we performed single molecule fluorescence *in situ* hybridization (smFISH) against Chrimson to directly measure its mRNA expression in the strains carrying the *syIs503[15xUAS::chrimson]* effector (Figure 1I). The results showed that for the strains with reconstituted cGAL in one TRN subtype, Chrimson level was comparable between the ALM and AVM strains and moderately lower in the PLM strain (Figure 1J). For the strains with Chrimson in two TRN subtypes, the Chrimson expression was similar between the two subtypes, although the AVM-PLM strain had generally stronger expression than the ALM-PLM strain (Figure 1J). Importantly, the highly correlated expression of Chrimson between the two subtypes in the same animal (Figure 1K) allows similar levels of optogenetic activation in the sensory neurons, so that the integration of anterior and posterior stimulation can be better assessed at the circuit level.

### Whole-field illumination stimulates specific TRN subtypes and generates varying responses

Using the Chrimson-expressing strains, we first applied whole-field illumination to activate specific TRN subtypes and measured the behavioral response of hundreds of animals (Figure 2A). Activation of only ALM or AVM resulted in the animal slowing down or reversing directions, which was reflected in the negative change of velocity, whereas activation of PLM led to acceleration or increase in velocity (Figure 2B). Using a cutoff of -0.05 mm/s in velocity change before and after the stimulus for slowing down or reversing and +0.05 mm/s for speeding up, we found that the activation of ALM or AVM results in significantly increased probability of slowdown or reversal and reduced probability of acceleration compared to no stimulation, whereas activation of PLM led to lower probability of slowing down and higher probability of speeding up (Figure 2B). These results are consistent with the expected behavioral outcome of activating the anterior or posterior touch response circuit only. The control strain carrying only *unkIs7[mec-17p::cGAL-N]* and *syIs503[15xUAS::chrimson]* did not show any change of velocity upon light stimulation. The baseline velocities of all Chrimson-expressing strains were similar (Figure S2A).

**Figure 2.**
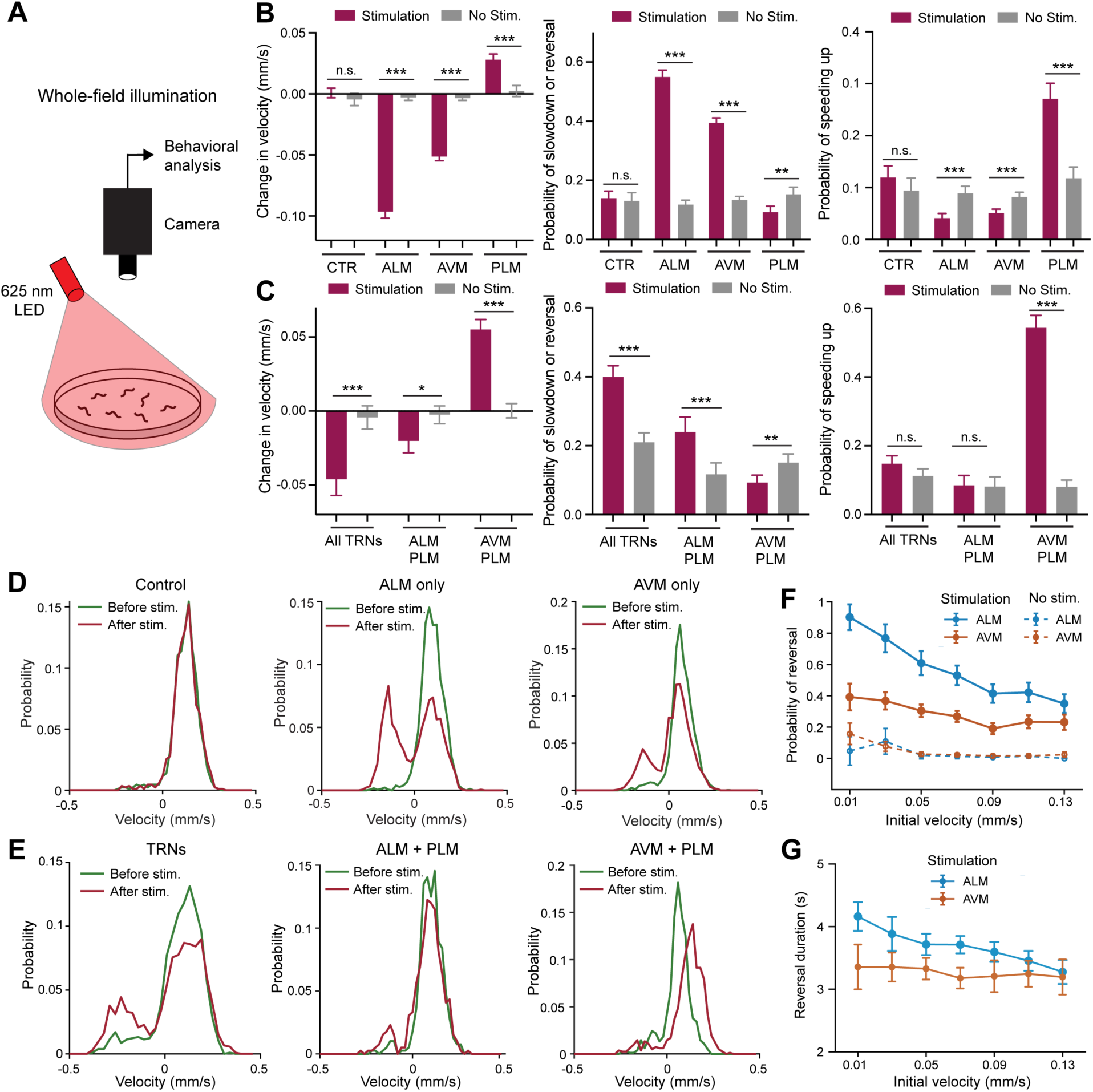
Whole-field illumination and behavioral analysis of animals with Chrimson expression in selective TRN subtypes. (A) Experimental setup for whole-field illumination. (B) Light-induced change in velocity, probability of slowdown/reversal (change in velocity ≤ -0.05 mm/s), and probability of speeding up (change in velocity ≥ 0.05 mm/s) of animals that express Chrimson selectively in one TRN subtype (≥ 680 stimulation events for each strain). For stimulation, red light with an intensity of 150 μW/mm^2^ was delivered; for non-stimulation, no light was delivered. The results are presented as mean ± 95% confidence interval in this Figure. One, two, and three asterisks indicate *p* < 0.05, 0.01, and 0.001, respectively, using Kolmogorov-Smirnov test with Bonferroni correction. The CGZ1409 *unkIs17; syIs503* strain that has no Chrimson expression in any TRNs was used as the control. (C) Light-induced change in velocity and probability of slowdown/reversal and speeding up for strains with multiple TRN subtypes expressing Chrimson (≥ 390 stimulation events for each strain). (D-E) Velocity distribution before and after the stimulation with 150 μW/mm^2^ red light in strains expressing Chrimson in no, one, or multiple TRN subtypes. (F) The relationship between the probability of reversal (defined by that velocity goes below -0.01 mm/s within 3 seconds of optogenetic stimulation) and the velocity at stimulus onset in the strains expressing Chrimson in ALM or AVM. (G) The relationship between reversal duration (defined as the period for which velocity is below -0.01 mm/s after stimulus onset) and initial velocity.

Next, we compared the response of ALM and AVM activation and found that activating the ALM neurons triggered a stronger anterior response than activating AVM, which is demonstrated by a bigger velocity change, higher probability of slowing down or reversing, and lower probability of speeding up (Figure 2B). When comparing the velocity distribution before and after the light stimulation, we found that ALM activation caused more animals to move backwards (having negative velocity) than AVM activation (Figure 2D and S2B). Moreover, the difference in behavioral response between ALM and AVM activation was more obvious when the animals were moving at a lower speed (Figure 2F-G). ALM activation was more likely to trigger reversal, and the reversals had longer duration when the velocity at stimulus onset is low compared to animals that were moving more quickly at stimulus onset. In contrast, AVM activation has a constantly lower probability of triggering reversal, and the reversals are shorter regardless of the initial velocity. These results suggest that the response of the animal to mechanosensory stimulation is often dependent on its locomotion state prior to the stimulation. Overall, the contribution of the two ALM neurons is stronger than that of the single AVM neuron in triggering backward movement upon anterior stimulations, which is consistent with previous finding that ablating the two ALM neurons produced more severe defects in anterior touch sensitivity than removing only the AVM neuron.^21^

Using the strains that expressed Chrimson in two TRN subtypes, we found that simultaneously activating ALM and PLM neurons resulted in a negative change of velocity and increased probability of slowdown or reversal, suggesting that the anterior response triggered by ALM activation outcompeted the posterior response triggered by PLM activation, so that the overall response is backward movement (Figure 2C). In contrast, simultaneous activation of AVM and PLM neurons led to forward acceleration, reduced probability of slowing down or reversing, and increased probability of speeding up, suggesting that the PLM-induced forward motion dominated over the AVM-induced backward motion (Figure 2C, 2E, and S2C). These results further support the idea that ALM neurons evoke stronger response in the downstream motor programs compared to AVM, possibly because optogenetic activation of the two ALM neurons constitutes a larger sensory input onto the interneurons that command motor output (Figure 1B) than activating the AVM neuron only.

Nonetheless, the sensory contribution of AVM to the circuit is still significant. By comparing the response of activating only ALM and PLM with the response of activating all TRNs (i.e., ALM, PLM, and AVM, since prior evidence suggests that PVM is not involved in the sensory-motor circuit^9^), we found that the additional activation of AVM led to more negative changes in velocity, higher probability of slowdown and reversal, and bigger difference between pre- and post-stimulation velocities (Figure 2C and 2E). These results and previous evidence^9,13^ reaffirm a sensory function of the AVM neuron.

### Targeted illumination of anterior and posterior TRNs reveals non-linear components of the touch circuit when integrating conflicting sensory information

For the strains expressing Chrimson in two or more TRN subtypes, the behavioral output under the whole-field illumination is the result of integrating the input of the anterior and posterior sensory neurons. To further investigate this signal integration, we used a previously developed close-loop targeted illumination system^14^ to deliver light with varying intensities to the head or tail of moving animals (Figure 3A and Movie S1). For the three strains (Chrimson in all TRNs, ALM-PLM, and AVM-PLM), head-only illumination led to deceleration or reversal, whereas tail-only illumination led to forward acceleration as expected (Figure 3B and C). Interestingly, when light was delivered to both head and tail, the backward locomotion triggered by anterior stimulation was countered to different extents by posterior stimulation in different strains. For example, when the head was stimulated at 40 μW/mm^2^ and the tail at 80 μW/mm^2^, we found that the strain with all TRNs expressing Chrimson still showed negative velocity change indicating deceleration, while the ALM-PLM strain had no obvious change in velocity, and AVM-PLM strain showed forward acceleration. These results suggested that when both ALM and AVM were activated, the tendency to move backward is strong and could not be countered even by posterior stimulation from illumination with light of twice the intensity of the anterior stimulation. If only ALM was activated, stronger posterior stimulation could cancel out the anterior stimulation; if only AVM was activated, the stronger posterior stimulation could outcompete the anterior stimulation to speed up the animal. These results support a model in which AVM activation alone evokes a weaker anterior response than the activation of the two ALM neurons, which is in turn weaker than activation all three anterior sensors.

**Figure 3.**
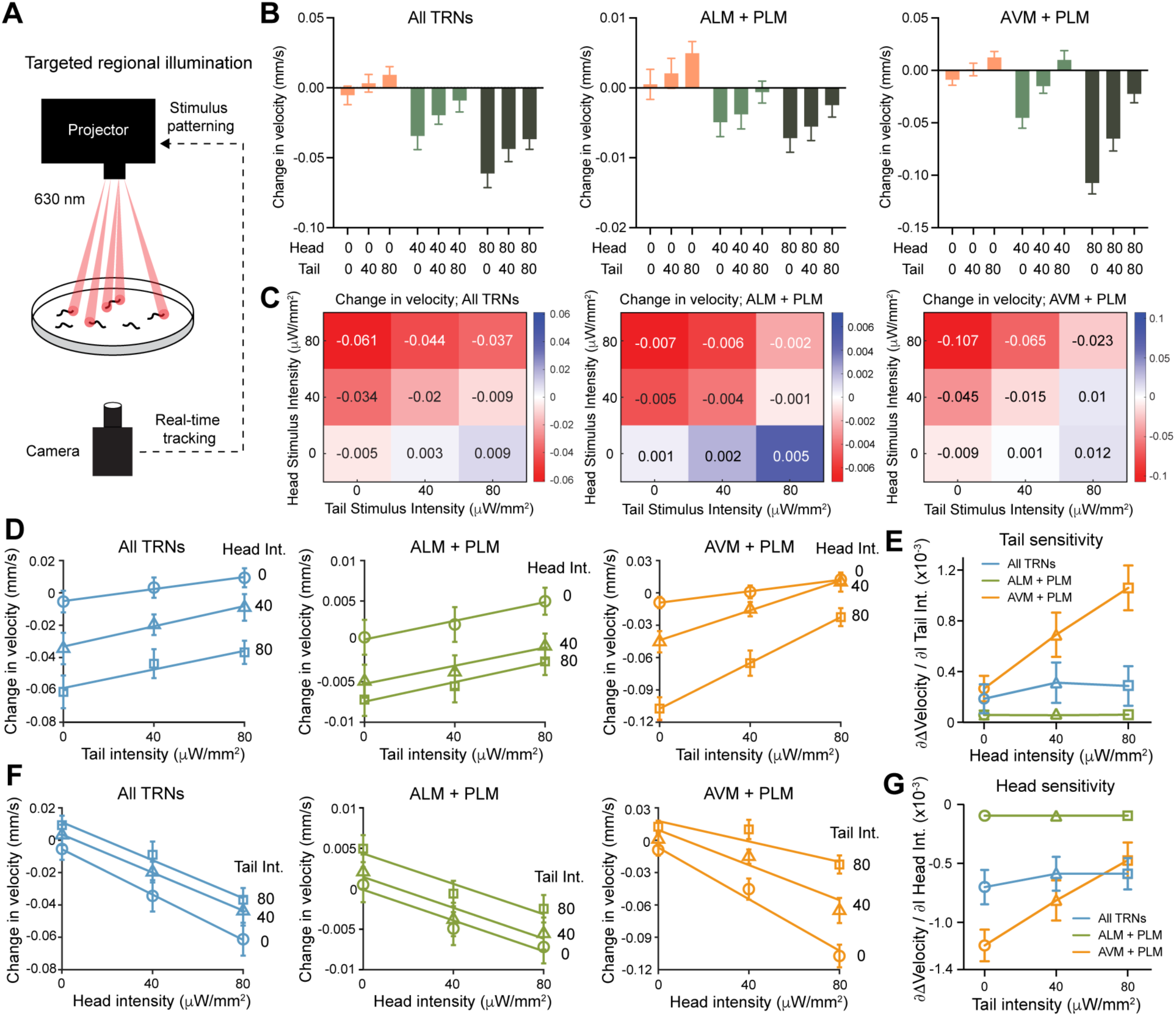
Targeted illumination reveals non-linear components of the touch circuit in the integration of conflicting sensory signals. (A) Experimental setup for targeted illumination of only head or tail regions of the animal. (B) Change in velocity under various conflicting sensory stimuli for the three strains expressing Chrimson in multiple TRN subtypes (≥ 597 stimulation events for each condition). Error bars show 95% confidence interval. (C) Mean velocity changes under various stimulation conditions are presented in a heatmap. (D) Change in velocity for a fixed head stimulus intensity but varying tail stimulus intensities. The line of the best fit is shown. (E) Tail sensitivity reports the slopes of the change-in-velocity curves against tail stimulus intensity, shown in (D), for each head stimulus intensity. (F) Change in velocity for a fixed tail stimulus intensity but varying head stimulus intensities shown as the line of the best fit. (G) Head sensitivity reports the slopes of the change-in-velocity curves against head stimulus intensity, shown in (F), for each tail stimulus intensity.

In addition to changes in velocity, the probability of head stimulus-induced slowing down or reversing was reduced more strongly by tail stimulation in the AVM-PLM strain compared to the all-TRN strain, while the probability of speeding up was also increased by a larger extent in AVM-PLM strain (Figure S3A-B). The ALM-PLM strain has generally weaker response to the same light stimuli than the other two strains due to lower Chrimson expression levels (Figure 1J), so that the effects on the probability of deceleration and acceleration (defined by ± 0.05 mm/s in velocity change before and after stimulation) were not very clear in the strain. However, we could still compare the antagonism between the anterior and posterior stimulation across strains in relative velocity changes normalized against the maximum change upon head- or tail-only stimulation for a given stimulus intensity (Figure S3C-F). For example, tail stimulation counters the head stimulation more strongly in the ALM-PLM strain compared to the all-TRN strain, whereas head stimulation counters the tail stimulation better in the all-TRN strain because of the additional activation of AVM neuron.

Next, we quantified how the velocity change responded to increasing tail (or head) stimulus intensities for a given head (or tail) stimulus. Interestingly, the all-TRN and the ALM-PLM strains responded similarly to the increasing intensity of tail (or head) stimulus regardless of the varying intensities of the head (or tail) stimulus, suggesting that the amplitude of the response can be founded by linearly combining the strength of local stimulation (Figure 3D-G). In other words, for these strains the combined response behaved as a linear contribution of what one would expect from summing the expected anterior and posterior response. In contrast, the AVM-PLM strain was more responsive to changes in the intensity of the tail stimulation when the head received a stronger stimulus (Figure 3D right panel) and conversely was less responsive to changes in the head stimulus intensity when the tail received a stronger stimulus (Figure 3F right panel). This is captured in the animal’s sensitivity to tail (or head) stimulation [called tail (or head) sensitivity in Figure 3E and 3G], which is calculated from the change in the slope of the plots of the animal’s response versus tail (or head) stimulus, for increasing head (or tail) stimulus intensities (Figure 3D and 3F).

We found that the tail (or head) sensitivity stayed constant across different intensities of head (or tail) stimulus for the all-TRN and ALM-PLM strains (constant slopes in Figure 3D and 3F and flat line in Figure 3E and 3G), which is expected for a linear system in which anterior and posterior responses are summed together. In contrast, the AVM-PLM strain had an increasing tail sensitivity with stronger head stimulation (the slope of velocity change became increasingly steep in Figure 3D right panel and the slope value increased in Figure 3E) and conversely a diminishing head sensitivity with elevating intensities of tail stimulus (the slope became increasingly flat in Figure 3F right panel and the absolute value of the slope decreased in Figure 3G). Thus, for the AVM-PLM strain, we observed non-linear integration of the anterior and posterior stimuli, suggesting that the contributions of anterior and posterior stimuli interact in a non-trivial and non-linear way. This cross-regulation between the anterior and posterior circuits is unexpected and, notably, was not visible when all TRNs are activated.

Taken together, our results suggest that when integrating the mechanosensory inputs from the anterior and posterior sensors, ALM and PLM neurons each contribute non-linear components to the overall touch response circuit that are strikingly balanced. When both ALM and PLM are activated, these non-linear contributions conspire to cancel out each other and the animal acts as if it combines anterior and posterior responses linearly, as evidenced by the ability of the ALM-PLM and the all-TRN strains to preserve a constant or near-constant sensitivity to head or tail stimuli. However, when only AVM but not ALM is activated at the anterior (in the AVM-PLM strain), the circuit balance required for linear output is disrupted, resulting in non-linear integration of the anterior and posterior signals. One way to visualize this non-linearity is to observe the warping of the surface fitted to the velocity change landscape under various stimulus conditions in the AVM-PLM strain (Movie S2, linear responses would have a flat surface without warping). Moreover, we reason that AVM’s contribution is primarily linear because, compared to the all-TRN strain, the ALM-PLM strain (in which AVM is not activated) still preserved the linear integration and exhibited constant head and tail sensitivities (Figure 3E and 3G). Therefore, we conclude that ALM and PLM individually make non-linear contributions to sensory integration, but when combined they work to preserve linear integration and sensitivity across a range of stimulus intensities.

### Whole-field illumination confirms stimulus intensity-dependent directional bias of AVM-PLM response predicted by the non-linearity of the circuit

The non-linear response of the AVM-PLM strain revealed an interesting relation of how sensitivity to head (or tail) stimulation changes with stimulation at the tail (or head) – the opposite side. It also makes several striking predictions. As the tail stimulation intensity increases (starting from zero), head sensitivity changes from being large and negative towards zero and, if this assumed linear relation holds, it predicts that eventually the head sensitivity could become positive by extrapolation (Figure 4A; in this case, positive number means that stronger head stimulation results in weaker response because the response is measured in negative velocity changes). Moreover, for a given head stimulus, we also observed a simple relation in this strain’s response to tail stimulus, as the velocity change seemed to increase linearly with tail intensities (Figure 3D and 4B). The above two linear relations allowed us to extrapolate from our targeted illumination data and make predictions on how the animal should responds to very large intensity stimulations if these trends were maintained (Figure 4C). Most intriguingly, we predicted that the animals would switch from slowdown or reversal to forward acceleration in response to equal head and tail illumination at sufficiently high intensities (Figure 4D).

**Figure 4.**
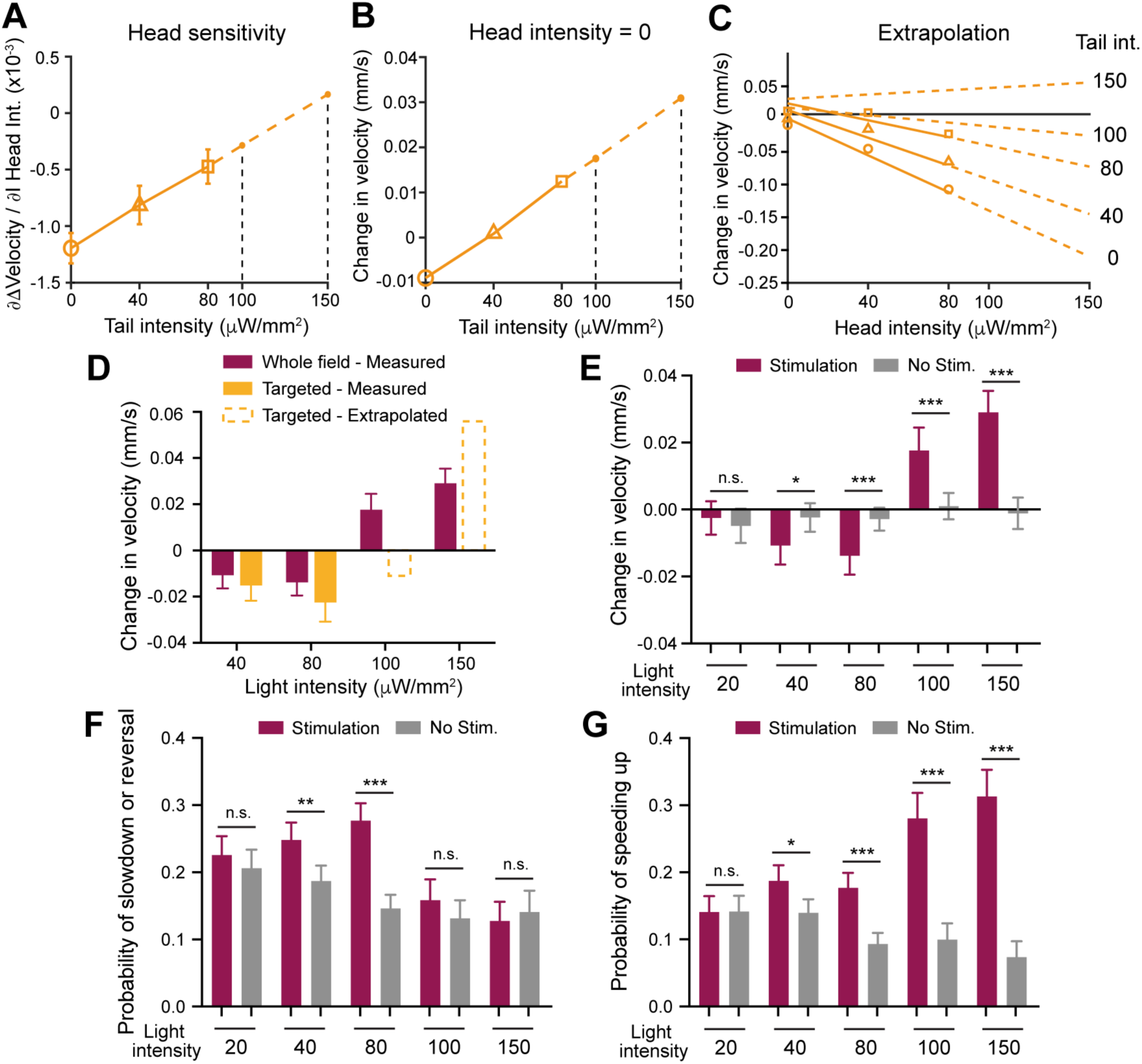
Non-linear integration of opposing signals in the touch circuit predicts stimulus intensity-dependent directional bias in the response of AVM-PLM strain. (A) Extrapolation of the head sensitivity (Figure 3G) for 100 and 150 μW/mm^2^ tail stimuli (dashed lines) for AVM-PLM strain. (B) Extrapolation of the expected change in velocity for head intensity of 0 μW/mm^2^ and tail intensities of 100 and 150 μW/mm^2^ (from Figure 3D right panel). (C) Change in velocity for a fixed tail intensity and varying head intensities (similar to Figure 3F right panel). The dashed lines represent extrapolated data determined in (A) and (B). (D) Comparison of targeted illumination responses (measured or predicted) to whole-field illumination responses for various light intensities. (E) Velocity change of the AVM-PLM strain under whole-field illumination as in (D) but also showing no stimulation controls (≥ 600 stimulation events for each condition). (F-G) Probabilities of slowdown or reversal (velocity change ≤ -0.05 mm/s) and of speeding up (velocity change ≥ 0.05 mm/s) for the AVM-PLM strain under various illumination conditions. Error bars in this Figure represents 95% confidence intervals.

To test this prediction, we used the whole-field illumination setup to activate both AVM and PLM with a range of light intensities from 20 to 150 μW/mm^2^. We found that at the highest intensities the measurements matched our predictions, as illuminating both the head and tail at 100 μW/mm^2^ indeed induced forward acceleration (Figure 4E). The threshold from a response of slowdown or reversal to forward acceleration appeared to be somewhere between 80 and 100 μW/mm^2^. The velocity distribution before and after the stimulus confirmed that the direction and magnitude of velocity change was dependent on the stimulus strength (Figure S4A). Similar stimulus intensity-based directional bias was not observed in the ALM-PLM strain and the all-TRN strain, since their response to the whole-field illumination at 150 μW/mm^2^ and targeted head-tail simultaneous illumination at 80 μW/mm^2^ were similar (i.e., deceleration or reversal) (Figure 2C, 2E, and 3B-C). These finding suggests that in the physiological condition, adding ALM neurons into the circuit provides a counteracting non-linear contribution to ensure a constant bias towards backward locomotion when integrating opposing signals at any given stimulus intensity.

## DISCUSSION

In this study, we combined the use of transgenic animals that express optogenetic actuators in specific neuronal subtypes of a circuit (through a genetic “AND” gate) with spatiotemporally precise delivery of light and high-throughput analysis of behavioral response. This approach allowed us to manipulate the activity of neurons at a single-cell resolution and elucidate their contribution to behaviors with high statistical power. A common technical concern of optogenetic manipulation is whether the light-induced activation of neurons operates within a physiologically relevant range. As a result, it remains difficult to compare experiments across strains and studies. Our study serves as one of the few examples that connect the expression level of the optogenetic actuators and light intensity with behavioral response in a quantitative manner. We found that under the same light stimulus, the extent of motor response scales almost linearly with the expression level of Chrimson (Figure S4B), while increasing light intensity also enhanced the strength of the response in a near-linear fashion (Figure 3D-G and 4B). Thus, the light stimulation we used was not likely to activate neurons at a saturating level, and we suspect that this may be the case for most optogenetic studies. Nonetheless, we advocate for the calibration of behavioral response against the expression levels of actuators and light intensity.

Being able to deliver light stimuli to specific TRN subtypes led us to uncover several new insights about the touch response circuit in *C. elegans*. First, previous studies found that when all TRNs are activated by plate tapping^22^ or optogenetic activation^23^, adult animals respond mostly by reversals. One hypothesis is that the three anterior sensory neurons (ALML, ALMR, and AVM) outcompeted the two posterior sensory neurons (PLML and PLMR) due to the activation of AVM as the tie breaker. Our results, however, suggest that even when only the two ALMs are activated, the anterior response could still outcompete the posterior response triggered by the two PLMs, suggesting that when conflicting signals were received, the touch circuit is intrinsically biased towards backward movement even if the postembryonic AVM is not directly activated. Nevertheless, further activation of AVM strengthens this bias towards deceleration or reversal.

Second, the fact that ALM-initiated backward locomotion can overpower PLM-initiated forward acceleration in the ALM-PLM strain suggests that the posterior circuit may have a generally lower efficacy than the anterior circuit, which may result from the different electrophysiological properties of the sensors. In fact, previous studies found that the mechanoreceptor current (MRC) in response to saturated stimuli was smaller in PLM compared to ALM neurons (54 versus 89 pA)^24,25^ and the Calcium response to the same mechanical stimulus was also smaller in PLM.^26^ Our data suggest that even when mechanoreception is bypassed by optogenetic activation, PLM may still be less effective than ALM neurons in triggering motor response either because PLM has a higher threshold for depolarization or because PLM is less efficient in activating relevant downstream interneurons.

A third insight is that when integrating opposing signals, the overall touch circuit contains antagonistic non-linear contributions from both ALM and PLM that conspire to generate a linear response to anterior or posterior stimuli across different intensities. This is most evident in the non-linear integration of head and tail stimulation in the AVM-PLM strain, which omits contributions from ALM and therefore reveals the non-linearity of PLM’s contribution. From the circuit connectivity (Figure 1B), PLM makes inhibitory chemical synapses with AVD interneuron that promotes backward movement,^21,27,28^ so PLM activation may induce AVD hyperpolarization in a non-linear fashion, which cannot be overcome by AVM-induced AVD activation. However, ALM activation may excite AVD non-linearly to overcome the inhibition from PLM and restore linear response to local stimuli. In addition, AVM (and not ALM) makes chemical synapses to the AVB-AVA pathway that controls the switch between forward and backward motor states.^21,27–29^ This connectivity may render AVM’s role in the circuit different from ALM’s and allow AVM to make linear contribution in the integration of conflicting signals.

The linear and non-linear components of the touch response circuit might serve different functions and thus maximize adaptability against changing environments. When all TRNs are activated, the response is linear, which allows accurate detection of the mechanical stimuli and evokes a response proportional to the intensity of the stimuli. Strikingly, this linearity in circuit response appears to be comprised of antagonistic non-linear contributions from the major sensors (i.e., ALM and PLM). Non-linear networks have mostly remained as theoretical systems for modeling complex neural circuits,^30–33^ while our work suggest that non-linearity can exist in a simple sensory reflex circuit with only a few neurons.

## Supporting information

Figure S1-S4 and Table S1; legends for Movie S1 and S2

## ACKNOWLEDGEMENT

We thank Hui Yuan Lim, Haoming He, and Ho Ming Terence Lee from the Zheng lab and Matthew Creamer from the Leifer lab for technical assistance and discussion. C.Z. is supported by funds from the National Natural Science Foundation of China (Excellent Young Scientists Fund for Hong Kong and Macau, 32122002), the Research Grant Council of Hong Kong (GRF 17105523, GRF 17106322, GRF 17113324, and CRF C7026-20G), and the Health Bureau of Hong Kong (HMRF 09201426). A.M.L is supported by the National Science Foundation through an NSF CAREER Award (IOS-1845137) and the Simons Foundation award SCGB 543003. Some strains used in this study were provided by the Caenorhabditis Genetics Center, which is funded by the NIH Office of Research Infrastructure Programs (P40 OD010440).

## Author contributions

C.Z. and A.M.L. conceived the study. L.T. and H.P. screened the promoters and created and characterized the transgenic strains. S.K. carried out the optogenetic stimulation experiments and analyzed the behavioral data. C.Z., S.K., and A.M.L prepared the manuscript with the input from L.T. C.Z. and A.M.L secured funding and supervised the project. All authors read and approved the manuscript.

## STAR ★ Methods

### KEY RESOURCE TABLE

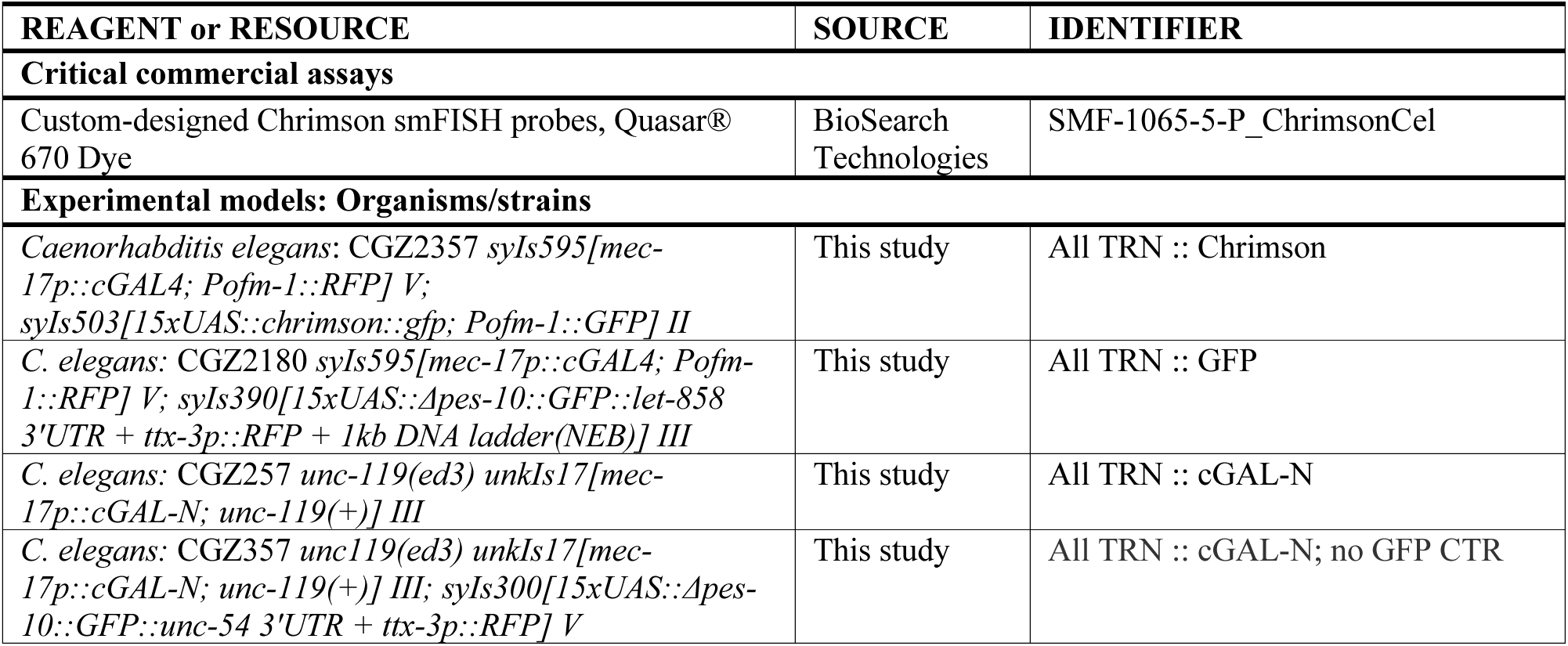
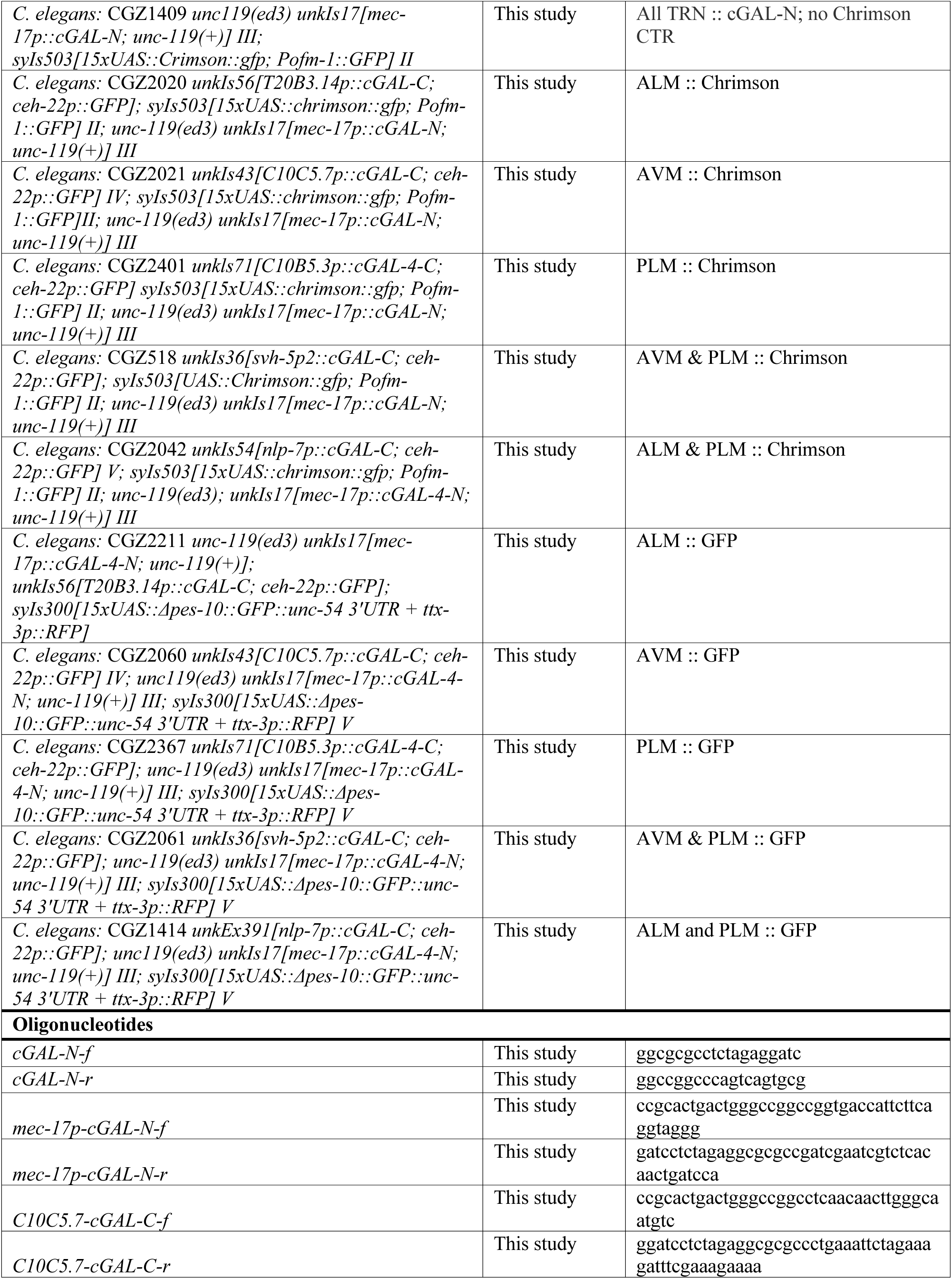
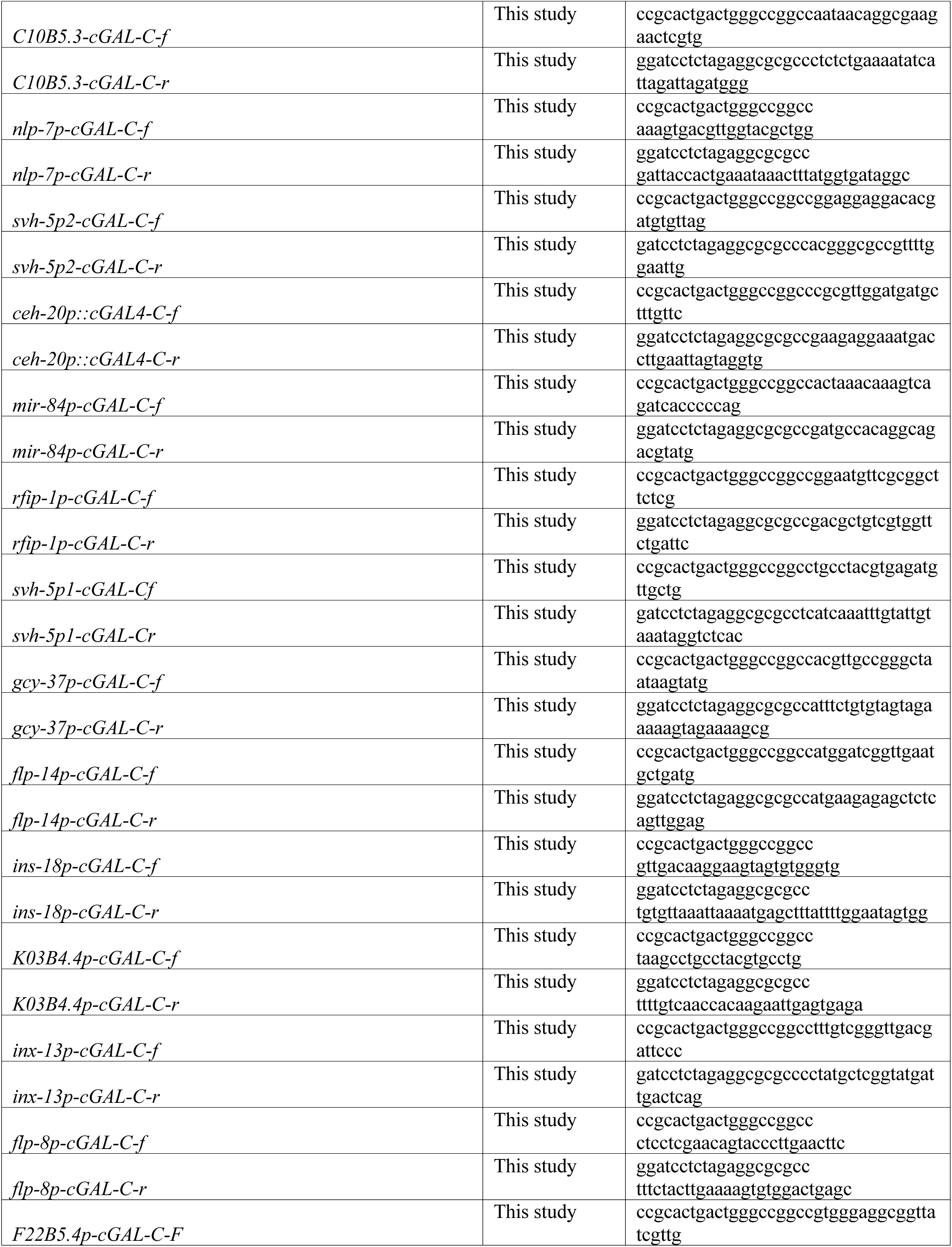
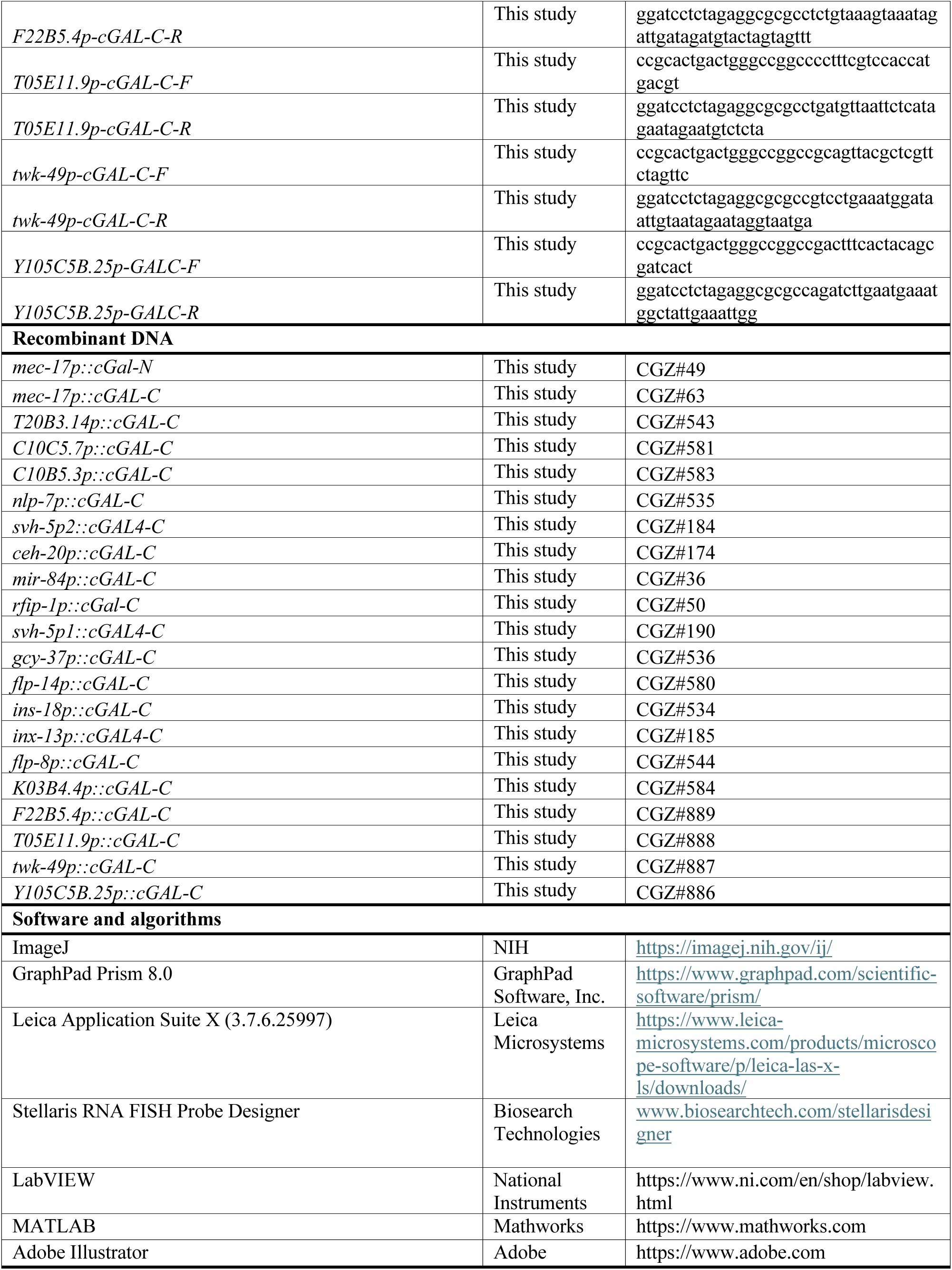

## RESOURCE AVAILABILITY

### Lead contact

Further information and requests for resources and reagents should be directed to and will be fulfilled by the lead contact, Chaogu Zheng.

### Materials availability

Strains and plasmids generated in this study will be shared by the lead contact upon request.

### Data and code availability

1. The raw behavioral data have been deposited to Figshare with the following DOI: 10.6084/m9.figshare.28217246 and are publicly available.
2. The code used to collect and analyze the behavioral dataset is available at https://github.com/leiferlab/kumar-touch-neuron-subtypes.git.
3. Any additional information required to reanalyze the data reported in this paper is available from the lead contact upon request.

## EXPERIMENTAL MODEL AND STUDY PARTICIPANT DETAILS

### C. elegans strains

*C. elegans* wild-type (N2) and transgenic strains were maintained as previously described.^34^ Most of the experiments were performed at 20°C on NGM plates seeded with *E. coli* (OP50) as food source unless otherwise indicated. The TRN-specific cGAL4 driver strain PS8373 *syIs574[mec-17p::cGAL4; Pofm-1::RFP]* and the effector strains PS8023 *syIs503[15xUAS::chrimson::gfp::let-858 3’UTR; Pofm-1::GFP]* and PS6843 *syIs300[15xUAS::Δpes-10::GFP::unc-54 3’UTR; ttx-3p::RFP]* were created by previous studies^6,16^ and obtained from the Caenorhabditis Genetics Center. The strains created in this study and used for behavioural studies are listed in the Key Resource Table.

## METHOD DETAILS

### DNA constructs and transgenesis

The *mec-17p::cGAL-N* plasmid was constructed by amplifying the cGAL-N coding sequence from pHW530 obtained from Addgene (#107131) and inserting the fragment between a 1.9 kb *mec-17* promoter and *unc-54* 3’UTR on a pDEST (Thermo Fisher) backbone using Gibson Assembly (New England Biolabs). This construct was injected together with *Cb_unc-119(+)* into *unc-119(ed3)* animals to make transgenic animals, which was then integrated by γ-irradiation, and the integrated line was outcrossed to make the strain CGZ257 *unc-119(ed3); unkIs17[mec-17p::cGAL-N; unc-119(+)]*. We then crossed CGZ257 with the effector strain PS6843 *syIs300 [15xUAS::Δpes-10::GFP::unc-54 3’UTR; ttx-3p::RFP] V* to create the strain CGZ357 *unc119(ed3); unkIs17; syIs300*, which is then used to test the possible intersectional expression of GFP in selective TRN subtypes.

To create the constructs that express cGAL-C from candidate promoters, the promoter sequences (e.g., a 3873-bp *T20B3.14* promoter upstream of the start codon; see Table S1 for details) were amplified from the genomic DNA and used to replace the *rab-3* promoter in the pHW522 [*rab-3p::cGAL-C::let-858 3’UTR*] plasmid obtained from Addgene (#107130). The resulted constructs were then injected together with *ceh-22p::GFP* (co-injection marker that expresses GFP in the pharynx) into CGZ357 to establish extrachromosomal arrays and to check for TRN subtype expression. The desired lines were then crossed with PS8023 *syIs503[15xUAS::chrimson::gfp::let-858 3’UTR, Pofm-1::gfp] II* to create lines that expressed Chrimson in specific TRN subtypes and to remove *syIs300* (the GFP signal from *syIs503* is very weak compared to *syIs300*). We then integrated the transgene using γ-irradiation and outcrossed the integrated lines six times. The integrated lines were crossed with CGZ357 again to reintroduce *syIs300* for the measurement of GFP intensity in the TRN subtypes. Strains, primers, and DNA constructs used in this study are listed in the Key Resource Table.

### Fluorescence imaging and single-molecule fluorescence in situ hybridization (smFISH)

Fluorescent imaging was performed on a Leica DMi8 inverted microscope equipped with a Leica K5 monochrome camera and the Leica THUNDER deconvolution system. Measurements of fluorescent intensity were made using the Leica Application Suite X (3.7.6.25997) software.

To measure the level of Chrimson mRNA in the TRNs, we designed and synthesized Stellaris FISH Probes against the Chrimson transcript using the Stellaris RNA FISH Probe Designer (www.biosearchtech.com/stellarisdesigner) from Biosearch Technologies (Petaluma, CA). Using a previous protocol,^35^ L4 animals of various strains were fixed, stained with the FISH probe labelled with Cy5 in the hybridization buffer, and then washed using the wash buffer (Biosearch Technologies). Hybridized animals were then mounted using VECTASHIELD Antifade Mounting Medium (Vector Laboratories #H-1000) and imaged using Leica DM8 with Cy5 filter sets. Single mRNA transcripts cannot be easily resolved due to the high expression level of Chrimson. So, the fluorescent intensity of the FISH staining in the cell bodies of TRN subtypes were quantified.

### Nematode handling for optogenetic stimulation

The following strains are used in the high-throughput behavioral analysis upon optogenetic stimulation: CGZ1409 (negative control), CGZ2357 (all TRN expressing Chrimson), CGZ2020 (ALM-specific Chrimson), CGZ2021 (AVM only), CGZ2401 (PLM only), CGZ518 (AVM and PLM), CGZ2042 (ALM and PLM). Details can be found in the Key Resource Table. All experiments were performed on day-one young adult animals. To obtain age synchronized young adults, gravid worms were bleached 3 days prior to the experiments. Bleached eggs were washed, collected, and resuspended in M9 and incubated on a shaker overnight. The following morning hatched L1 larvae were centrifuged and transferred to freshly seeded plates consisting of 1 ml of 0.5 mM all-trans-retinal mixed with OP50 and stored in the dark at 20°C until young adulthood. For experiments, young adult worms were washed in M9 and transferred to an empty agarose plate for experiments. Excess M9 solution was absorbed with a Kim wipe as described previously.^36^

### Whole-field optogenetic stimulation assay

To deliver red light to the entire animal, we used a previously published whole-field optogenetic delivery system.^23^ A plain agarose plate containing 40-50 freely moving worms was illuminated by an 850 nm Infrared LED. For imaging the worms, a 2592 × 1944 pixel CMOS camera (ACA2500-14um, Basler) was used to capture image of the plate at 14 frames per second and a magnification of 20 μm per pixel. Optogenetic stimulation was delivered using three 625 nm LEDs (M625L3, Thorlabs) placed in a way that their light uniformly illuminated the camera’s field of view on the agarose plate. A 3 sec light pulse with an inter-trigger interval of 30 sec was delivered to the agarose plates on which worms are moving freely. Two light intensities were randomly delivered to the worms: 150 μW/mm^2^ (experiment) and 0 μW/mm^2^ (control). Specifically, for the AVM-PLM strain, we also tested other light intensities, including 20, 40, 80, and 100 μW/mm^2^ for the whole-field illumination setup, the results of which were shown in Figure 4E-G.

A post-processing algorithm was used to identify behaviors of worms through pose dynamics classification as previously described.^23,36^ Briefly, camera frames were analyzed to calculated various behavior parameters of each worm such as position, body curvature, centerline, etc. The first and last index of the centerline were first assigned as head and tail, respectively; we then observed the entire duration of the moving trajectory of the animal to make a more definitive assignment and correct any errors in assigning head and tail. Unhealthy worm with prolonged pauses were also identified and excluded from the analysis. The algorithm defined behavior of the worm by classifying pose dynamics into a behavioral map of forward, reverse, and turns.^14,23^

### Targeted head-tail optogenetic stimulation assay

To deliver red light to the head or tail of the worm to optogenetically activate the anterior and posterior mechanosensory neurons separately, we used a previously published high-throughput close-loop optogenetic delivery system.^14,36^ The system consists of three key components: a camera for imaging and tracking the worms, a projector to deliver optogenetic stimuli, and a plain agarose plate consisting of 40-50 freely moving worms. A 3 sec, 500-µm diameter red light stimulation of 630 nm was delivered to either the head, tail, or both simultaneously to each tracked worm on the agarose plate. The illumination spot was centered on the tip of the head or tail of the worm. The inter-trigger stimulus interval was 30 seconds. A stimulus with an intensity randomly chosen from the set of 0, 40, and 80 µW/mm^2^ was delivered to the head or tail or both of the worm. Thus, there were in total nine stimulus combinations: 1) Head: 0 µW/mm^2^, Tail 0 µW/mm^2^ 2) Head: 0 µW/mm^2^, Tail 40 µW/mm^2^, 3) Head: 0 µW/mm^2^, Tail 80 µW/mm^2^, 4) Head: 40 µW/mm^2^, Tail 0 µW/mm^2^ 5) Head: 40 µW/mm^2^, Tail 40 µW/mm^2^, 6) Head: 40 µW/mm^2^, Tail 80 µW/mm^2^, 7) Head: 80 µW/mm^2^, Tail 0 µW/mm^2^ 8) Head: 80 µW/mm^2^, Tail 40 µW/mm^2^, and 9) Head: 80 µW/mm^2^, Tail 80 µW/mm^2^.

In this targeted optogenetic stimulation setup, two independent sets of behavior mapping pipelines were used, one for real-time tracking of worms and optogenetic stimulus delivery, and the other for post-processing analysis.^14,36^ The real-time algorithm in LabVIEW tracked all the worms on the agarose plate and calculated various behavior parameters of each worm and assigned the head and tail based on the first and last index of the centerline, respectively. Every 30 seconds, the real-time algorithm signals the computer-controlled projector to deliver optogenetic stimulus to the head, tail, or both simultaneously for every tracked worm with one of the nine stimulus intensity combinations. The post-processing algorithm is similar to the one used in the whole-field optogenetic stimulation assay. Erroneous head-tail assignment was corrected by observing longer trajectory, unhealthy worms were excluded, and behaviors were assigned based on pose dynamics.

### Data analyses

To calculate the change in velocity, we only considered those worms that are moving forward at the stimulus onset. Hence, any worm with a negative velocity (moving backwards) at stimulus onset was discarded from further analysis. Change in velocity was defined as the difference between the velocity at the end of 3 sec optogenetic stimulus (*v_final*) and the velocity at the beginning of the optogenetic stimulus (*v_initial*). So, change in velocity= *v_final* - *v_initial*.

To calculate the probability of slowing down or reversal and the probability of speeding up, we defined a worm that is slowing down or reversing if the change in velocity of the worm upon optogenetic stimulation is lower than or equal to 0.05 mm/s. The binary array of 1 (worm that slowed down or underwent reversal) and 0 (worm that did not) was used to determine the probability. Similarly, the probability of speeding up was calculated by defining speeding up as change in velocity upon optogenetic stimulation to be greater than or equal to 0.05 mm/s.

To calculate the relationship between initial velocity and the probability of reversal, we only considered the forward moving animals and determined the initial velocity of the worms at the stimulus onset for all the stimulus events. The initial velocities were then grouped into bins of 0.02 mm/s increments. Thus, the centers of these bins are 0.1, 0.3, 0.5, 0.7, 0.9, 0.11, and 0.13 mm/s. A worm is considered to have reversed, if the velocity of the worm goes below a threshold of -0.01 mm/s during the three-second optogenetic stimulus window. The binary array of 1 (worm that reversed) and 0 (worm that did not) is used to determine the probability of reversal. This was repeated for all the worms in various bins to calculate the probability of optogenetically evoked reversal at different initial velocity.

## QUANTIFICATION AND STATISTICAL ANALYSIS

In our high-throughput behavioral analysis, stimulus events are the fundamental unit, and we reported the proportion of all stimulus events that result in various behavioral outcome, the total number of stimulus events, and the corresponding 95% confidence interval. Statistical significance for the difference between the stimulation and non-stimulation conditions were analyzed using the Kolmogorov-Smirnov test with Bonferroni correction. In Figure 3D-G, we used the MATLAB’s built-in functions “polyfit” to fit a polynomial of degree 1 for the line of best fit and used the “confint” function to determine the 95% confidence intervals.

