## Supplementary material for "Functional Interrogation of Neuronal Subtypes via Intersectional Expression of Optogenetic Actuator Reveals Non-linear Components in a Linear Circuit": Figure S1-S4 and Table S1; legends for Movie S1 and S2

**Supplementary Table, Movies, and Figures**

**Table S1. Promoters tested for TRN subtype-specific transgene expression, related to Figure 1.** Red ones are selected promoter combinations for split cGAL expression. “N.D.” means “not described” before.

**Movie S1. Examples of targeted illumination to optogenetically activate specific TRN subtypes, related to Figure 3.** Optogenetic stimulation was delivered to either the anterior or the posterior or both regions of the worm. Green dot denotes the worm’s head. Green line denotes the centerline. The yellow line shows the trajectory of a midpoint along the worm’s centerline over the past 10 seconds. Red region indicates area illuminated by red light. Left: optogenetic stimulation delivered to the head of the worm, middle: optogenetic stimulation delivered to the tail, right: optogenetic stimulation delivered to both the head and tail.

**Movie S2. Rotating surface plot for the velocity change of animals under various stimulus intensities, related to Figure 3.** Heatmap data shown in Figure 3C are fitted with a surface using the “surf” function of MATLAB for all the three strains. The AVM-PLM strain shows significant amount of warping compared to the all-TRN and the ALM-PLM strain.

**Figure S1. Selective labeling of TRN subtypes at various developmental stages, related to Figure 1.**

**Figure S2. Baseline velocity of the optogenetic strains, related to Figure 2.**

**Figure S3. Probability of slowdown/reversal and speeding up and relative percentage of velocity change under the targeted illumination setup, related to Figure 3.**

**Figure S4. AVM-PLM strain’s response under whole-field illumination with varying light intensities and the correlation between Chrimson level and behavioral response, related to Figure 4.**

**Table S1. Promoters tested for TRN subtype-specific transgene expression.** Red ones are selected promoter combinations for split cGAL expression. "N.D." means "not described" before.

| Promoter 1 for cGAL-N | Promoter 2 for cGAL-C | Promoter 2 length | Reporter expression for promoter 2 in TRNs | scRNA-seq expression for Promoter 2 (TPM) | | | | Observed expression pattern with UAS::GFP |
|---|---|---|---|---|---|---|---|---|
| | | | | ALM exp. | AVM exp. | PVM exp. | PLM exp. | |
| *mec-17* | - | | - | - | - | - | - | None |
| *mec-17* | *mec-17* | 1901 bp | ALM, AVM, PLM, PVM | 179723 | 87286 | 52678 | 161259 | ALM, AVM, PLM, PVM |
| *mec-17* | *T20B3.14* | 3873 bp | N.D. | 2687 | 0 | 0 | 0 | ALM only |
| *mec-17* | *C10C5.7* | 2033 bp | N.D. | 0 | 12367 | 0 | 0 | AVM only |
| *mec-17* | *C10B5.3* | 3945 bp | N.D. | 0 | 0 | 0 | 1539 | PLM only |
| *mec-17* | *nlp-7* | 2763 bp | ALM and PLM (PMID:19875417) | 2160 | 0 | 0 | 2251 | ALM and PLM |
| *mec-17* | *svh-5p2* | 4697 bp (*a* isoform) | PLM (PMID: 26539892) | 79 | 39 | 50 | 158 | AVM and PLM |
| *mec-17* | *ceh-20* | 2291 bp | ALM (PMID: 26547238) | 91 | 36 | 18 | 19 | ALM only |
| *mec-17* | *mir-84* | 1943 bp | ALM and AVM (PMID: 23599497; 26539892) | - | - | - | - | ALM, AVM, and PLM |
| *mec-17* | *rfip-1* | 2885 bp (*d* isoform) | PLM (PMID: 26539892) | 0 | 0 | 0 | 181 | ALM and PLM |
| *mec-17* | *svh-5p1* | 4245 bp (*b* isoform) | PLM (PMID: 26539892) | 79 | 39 | 50 | 158 | ALM, AVM, and PLM |
| *mec-17* | *gcy-37* | 1237 bp | AVM (PMID: 35226663) | 0 | 315 | 0 | 0 | None |
| *mec-17* | *flp-14* | 3892 bp | None (PMID: 15236235) | 0 | 38564 | 0 | 0 | None |
| *mec-17* | *ins-18* | 3454 bp | N.D. | 0 | 0 | 0 | 1699 | None |
| *mec-17* | *inx-13* | 1526 bp | PLM (PMID: 26539892) | 0 | 0 | 0 | 0 | None |
| *mec-17* | *flp-8* | 2613 bp | PVM (PMID: 15236235) | 0 | 12629 | 73381 | 0 | Variable AVM and PVM |
| *mec-17* | *twk-49* | 1443 bp | N.D. | 0 | 0 | 1802 | 0 | Variable PVM |
| *mec-17* | *K03B4.4* | 5323 bp | N.D. | 0 | 0 | 8964 | 0 | None |
| *mec-17* | *F22B5.4* | 5177 bp | N.D. | 0 | 0 | 2837 | 0 | None |
| *mec-17* | *T05E11.9* | 2590 bp | N.D. | 0 | 0 | 2413 | 0 | None |
| *mec-17* | *Y105C5B.25* | 4527 bp | N.D. | 0 | 0 | 1108 | 0 | None |

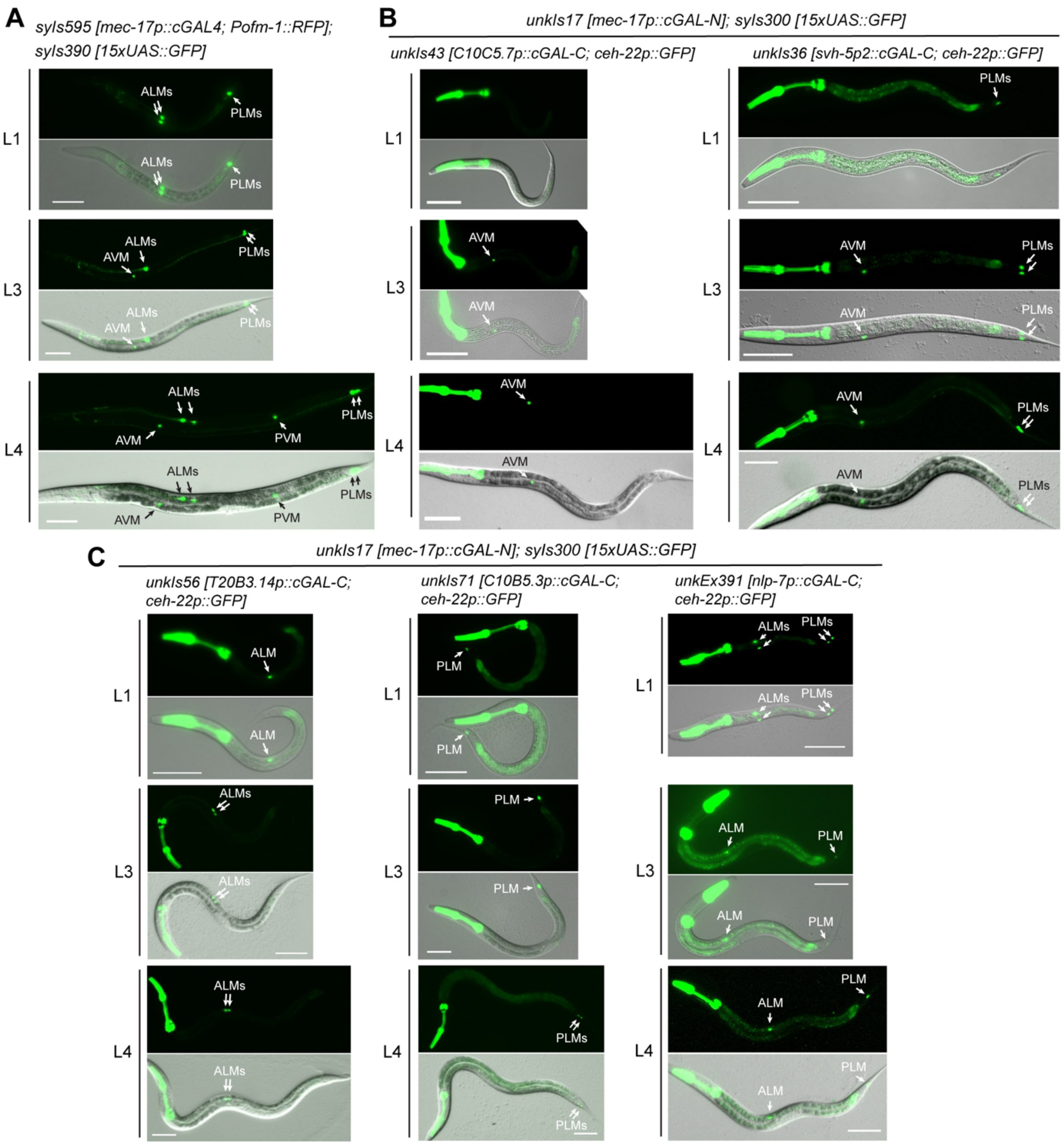


**Figure S1. Selective labeling of TRN subtypes at various developmental stages, related to Figure 1.** (A) GFP expression in CGZ2180 *syIs595[mec-17p::cGAL4]; syIs390[15xUAS::GFP]* animals at different larval stages. GFP in AVM was observed from L3 stage onward, and GFP in PVM was observed from L4 stage onward. (B) GFP expression in the CGZ2060 *unkIs17; syIs300; unkIs43* strain labeling AVM only and the CGZ2061 *unkIs17; syIs300; unkIs36* strain labeling both AVM and PLM. GFP in AVM was only visible from L3 stage onward (>19 hours post hatching). (C) GFP expression in strains that label ALM or PLM or both. GFP expressions were seen from L1 stage onward. Extrachromosomal array was used for *nlp-7p::cGAL-C*, since the integrated line was linked to *syIs300*. Scale bars = 100 μm.

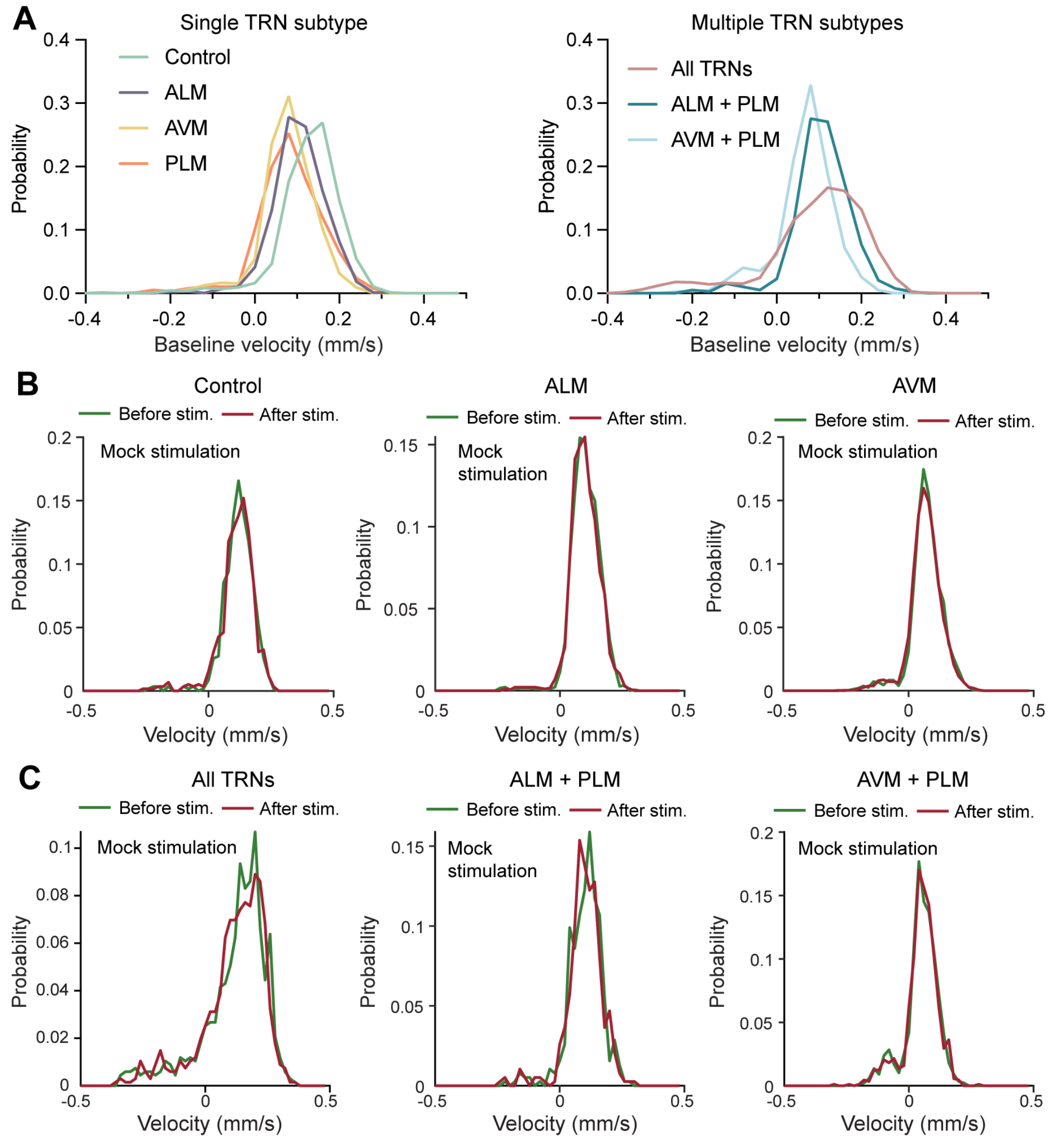


**Figure S2. Baseline velocity of the optogenetic strains, related to Figure 2.** (A) The distribution of baseline velocity of the animals expressing Chrimson in selective TRN subtypes. (B-C) The velocity distribution of various optogenetic strains before and after the mock stimulation (0 $\mu W/mm^2$) in the whole-field illumination experiment.

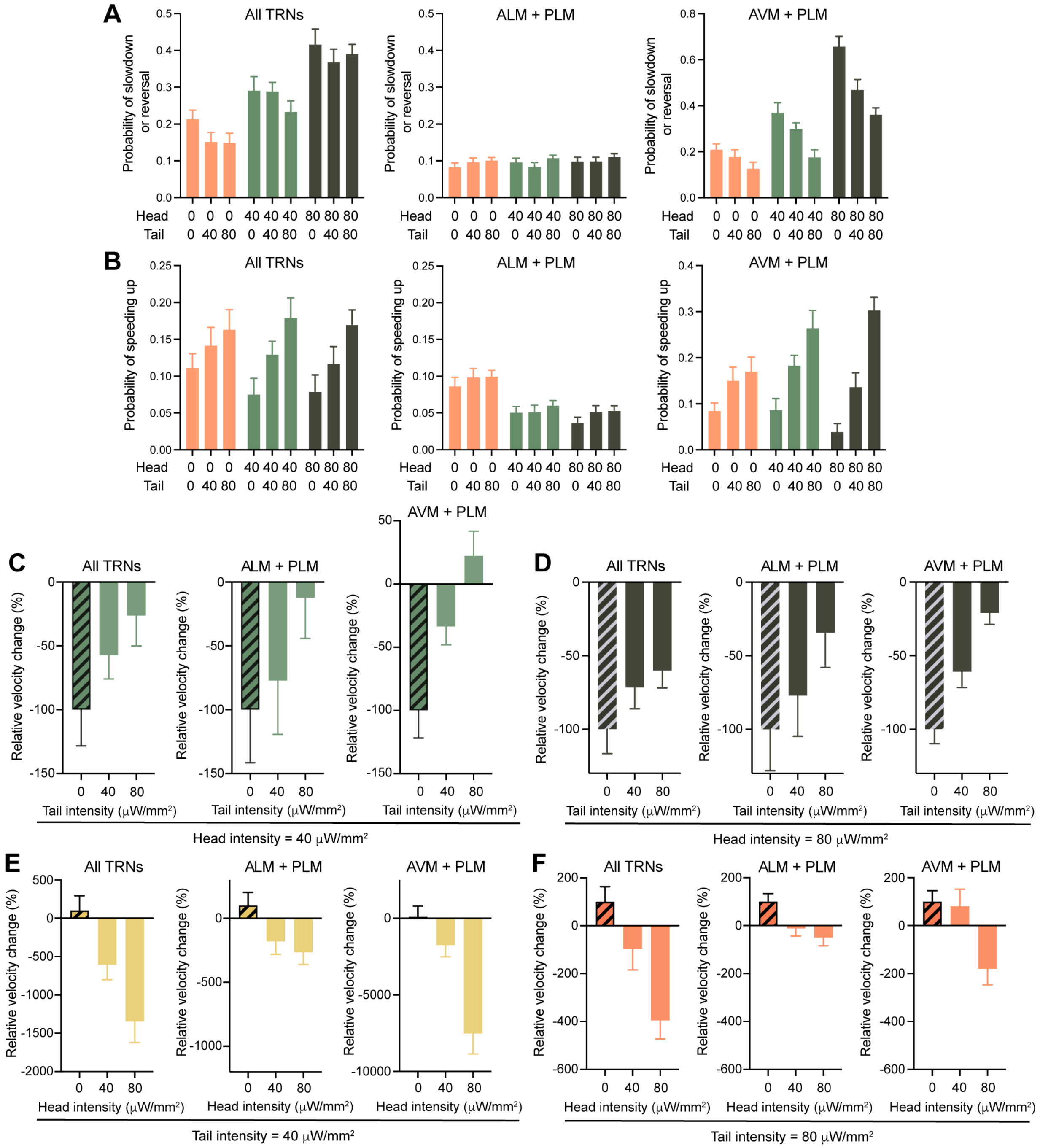


**Figure S3. Probability of slowdown/reversal and speeding up and relative percentage of velocity change under the targeted illumination setup, related to Figure 3.** (A-B) Probability of slowdown or reversal (velocity change ≤ -0.05 mm/s) and speeding up (velocity change ≥ 0.05 mm/s) of the strains expressing Chrimson in multiple TRN subtypes. Three light intensities (0, 40, and 80 μW/mm$^2$) for head and tail illumination were used in combinations. (C-D) The relative velocity change of the all-TRN, ALM-PLM, and AVM-PLM strains under the same head stimulus (intensity = 40 or 80 μW/mm$^2$) but varying tail stimulus (intensity = 0, 40, and 80 μW/mm²) with the absolute velocity change normalized to the velocity change upon head-only stimulation (i.e., head = 40 or 80 μW/mm²; tail = 0 μW/mm²). The directionality of the velocity change is preserved. Error bars indicate normalized 95% confidence interval. (C-D) Relative velocity change of the three strains under the same tail stimulus (intensity = 40 or 80 μW/mm²) but varying head stimulus. The absolute velocity change was normalized to the velocity change under tail-only stimulation (i.e., tail = 40 or 80 μW/mm²; head = 0 μW/mm²). AVM-PLM strain had very small response (0.00087 mm/s) under tail = 40 μW/mm² and head = 0 μW/mm². So, the relative velocity change showed very high percentage for tail = 40 and head = 40 or 80 μW/mm². For (C-F), the bar filled with diagonal stripes in each panel is used as reference to determine the relative percentage for other stimulus conditions.

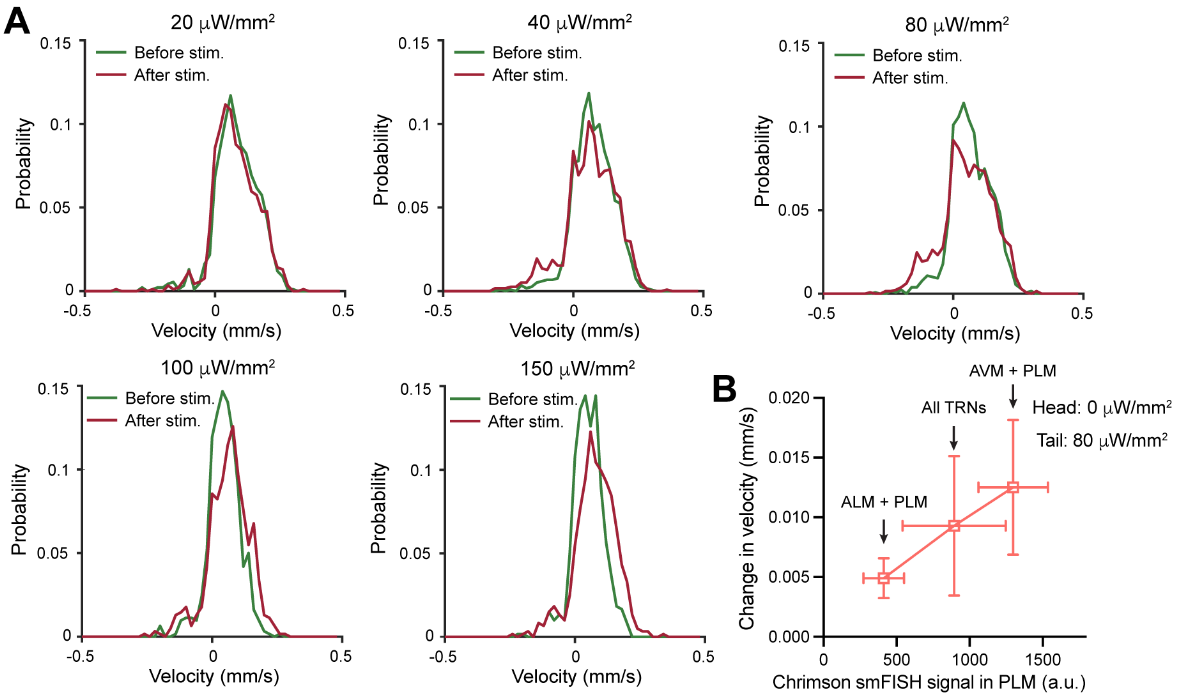


**Figure S4. AVM-PLM strain's response under whole-field illumination with varying light intensities and the correlation between Chrimson level and behavioral response, related to Figure 4.** (A) Velocity distribution of the AVM-PLM strain under various stimulus intensities in the whole-field illumination experiments related to Figure 4E-G. (B) Change in velocity (mean ± 95% confidence interval) were plotted against Chrimson smFISH intensity (mean ± SD) in the PLM neurons of the three strains when head intensity = 0 μW/mm² and tail intensity = 80 μW/mm².
